# Residue burial encodes a protein’s fold

**DOI:** 10.64898/2026.03.28.714986

**Authors:** Alex T. Grigas, Jacob Sumner, Corey S. O’Hern

## Abstract

Protein structure is controlled by a high-dimensional energy landscape, which is a function of all of the atomic coordinates of the protein. Can this landscape be accurately described by a low-dimensional representation? We find that residue core identity, a binary *N*-dimensional encoding indicating whether each of the *N* amino acids in a protein is buried in the core or not, can predict the protein’s backbone conformation more efficiently than all other representations that we tested. Core identity is 4 times more efficient than previous estimates of the bits per residue needed to encode a protein’s native fold, 2 times more efficient than the C_*α*_ contact map, and 1.5 times more efficient than the machine-learned embeddings from FoldSeek’s 3Di. Even when the folded structure is unavailable, predicting each residue’s burial from sequence yields a more accurate estimate of fold quality than predicting pairwise contacts from the same sequence information. Thus, this work emphasizes that the problem of determining a protein’s native fold can be re-framed as predicting each residue’s core identity.

---

How much information is needed to fold a protein? One possible answer is that we need enough information to constrain all of the degrees of freedom in the protein. There are three backbone dihedral angles per residue, but the dihedral angle across the peptide bond is typically constrained to be flat, leaving 2*N* backbone degrees of freedom for a protein with *N* residues. These backbone dihedral angles take on a restricted range of values in the Ramachandran plot due to intra-residue atomic clashes [1–3]. While these restricted angle ranges reduce the information needed to specify protein structure, they do not completely constrain the continuous dihedral angle degrees of freedom. Thus, identifying the minimal structural information needed to specify the native fold of a protein is still a fundamental open question.

Prior work has presented an intuitive physical picture of protein folding: proteins fold due to the hydrophobic effect, which drives hydrophobic collapse with hydrophobic residues buried in the solvent-inaccessible core, while polar and charged residues are exposed to solvent on the protein’s surface [4]. Analysis of core packing has yielded key insights into protein structure [5–13], yet until now core packing studies have not been able to accurately determine a protein’s native fold. In contrast, several end-to-end machine learning methods, which leverage large evolutionary datasets, can often predict protein structure accurately from sequence [14–18]. However, with these methods, it is unclear whether we have any deeper physical understanding of the protein folding process or even why these methods are so accurate.

In this Letter, we show that residue core identity, a binary label that identifies each residue as core versus non-core, is not only sufficient to determine whether computational models for protein structure are accurate, but it is also the most information-efficient encoding of protein structure among all of the physical and machine-learned representations that we tested. Previous work using thermodynamic and sequence alignment data suggested that 2-3 bits per residue of information are needed to accurately fold proteins [19]. However, we find that residue core identity encodes the native fold using only bits per residue, which is a fourfold improvement over the prior estimate, two times more efficient than the information in the full C_*α*_ contact matrix, and 1.5 times more efficient than the machine-learned representation from FoldSeek’s 3Di [20]. Further, residue core identity can be predicted directly from sequence and using it to encode protein structure is robust to labeling noise.

Our approach is to encode structural features of the native fold, such as the C_*α*_ contact map and each residue’s core identity, and determine the accuracy with which each encoding predicts the protein structure. In information-theoretic terms, each structural feature defines a channel, and we vary the size of the message sent through that channel. If a channel can predict the protein structure with high fidelity, it contains sufficient information to identify the native fold. Thus, maximizing the predicted accuracy over an ensemble of candidate structures would recover the native backbone conformation.

We begin by building a dataset of computationally-generated structural models of experimental high-quality x-ray crystal structures from the protein folding competitions CASP11-15, resulting in 63 targets and ∼24, 000 predicted structures. (See End Matter.) In CASP, research groups predict each target protein’s structure from its sequence, resulting in a dataset of diverse structural models of each protein that vary in accuracy compared to the targets. We use the average Local Distance Deviation Test (LDDT) to quantify the accuracy of the positions of the C_*α*_ atoms of the predicted structures compared to the targets [21]. 0 ≤ LDDT ≤ 1, where 1 indicates that the predicted structure matches the target structure.

Next, as an illustrative example, we show that the pairwise C_*α*_ contact map can be used to encode a protein’s fold. We define the contact map as

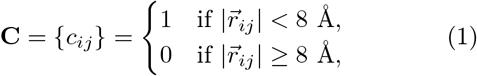

where 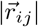 are the *N*(*N* − 1)/2 distances between C_*α*_ atoms on residues *i* and *j* and an 8Å distance cutoff is often used to filter the full distance matrix into binary contact labels. We denote **C**_*n*_ and **C**_*p*_ as the contact maps of the native fold and CASP models, respectively. (See Fig. 1 (a).)

**FIG. 1.**
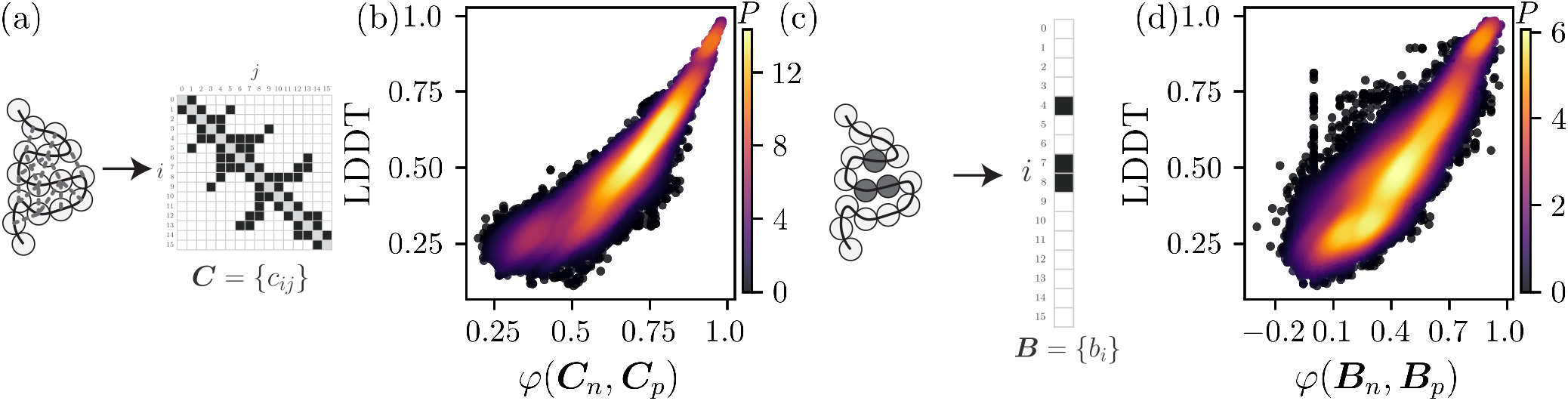
(a) A schematic of a folded protein represented by a polymer of connected disks (C_*α*_ atoms) with non-bonded pair-wise C_*α*_ contacts (grey dashed lines) encoded in the contact map **C** = {*c*_*ij*_}, where *c*_*ij*_ = 1 (black) represents a contact and 0 represents non-contact (white). (b) The probability density *P* (increasing from dark to light) plotted as a function of the accuracy of the backbone conformation LDDT and contact map similarity *φ*(**C**_*n*_, **C**_*p*_) between the native fold **C**_*n*_ and model **C**_*p*_. (c) Similar schematic of a folded protein to that in (a) with the core residues colored grey and represented by the core residue identity vector **B** = {*b*_*i*_ }, where *b*_*i*_ = 1 (black) represents a core residue and 0 (white) represents a non-core residue. (d) Probability density *P* (increasing from dark to light) plotted as a function of LDDT and the core identity similarity *φ*(**B**_*n*_, **B**_*p*_) between the native fold **B**_*n*_ and model **B**_*p*_.

To quantify how well a predicted structure’s contact map matches the native structure, we use Matthews correlation coefficient,

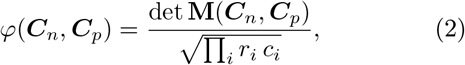

where **M**(***C***_*n*_, ***C***_*p*_) is the 2 × 2 confusion matrix with diagonal elements that count the number of matches between the ones (and separately between the zeros) of **C**_*n*_ and **C**_*p*_ and with off-diagonal elements that count the number of mismatches between the zeros and ones, and *r*_*i*_ and *c*_*i*_ are the sums of row and column *i* of **M** [22]. *φ* = 1 when **C**_*n*_ = **C**_*p*_ and *φ* = 0 when the predicted labels have no correlation to the labels of the native fold. In Fig. 1 (b), we plot the probability density for obtaining the similarity between the native and predicted C_*α*_ contact maps *φ*(**C**_*n*_, **C**_*p*_) and LDDT. As expected, we find a strong Spearman correlation *ρ* = 0.95 between *φ*(**C**_*n*_, **C**_*p*_) and LDDT.

How well does the residue core identity correlate with LDDT? We quantify residue burial using the relative solvent accessible surface area (rSASA) [23, 24]. rSASA determines what fraction of an amino acid’s surface area is exposed to a probe the size of a mater molecule. We then define residue burial as

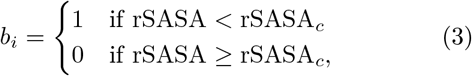

where b_*i*_ is the core identify of residue *i* and ***B*** is the complete set of *N* labels for the protein. (See Fig. 1 (c).) As with the contact map, we can score each predicted structure by calculating the similarity between the native and predicted core identity sets *φ*(***B***_*n*_, ***B***_*p*_). In Fig. 1 (d), we plot the probability density for obtaining *φ*(**B**_*n*_, **B**_*p*_) and LDDT and find a remarkably large Spearman correlation between *φ*(**B**_*n*_, **B**_*p*_) and LDDT of 0.94. We emphasize that the core identity label does not require that the core residues are near each other, just that rSASA < rSASA_*c*_. We show in Fig. 5 in the End Matter that these results are relatively insensitive to the choice of rSASA_*c*_.

Both pairwise contacts and core identity carry enough information to predict LDDT. However, to fairly compare the two metrics we need to consider the amount of information that they contain. For example, the C_*α*_ contact map is sparse, with many more non-contacts than contacts. Therefore, we not only need to compare the number of labels used, but also the information of each label *ι*(*x*) = − log_2_(*p*(*x*)), where the base two sets the units to be in bits of information and *p*(*x*) is the probability for obtaining outcome *x*. For the target x-ray crystal structures, contacts are much less frequent than non-contacts with *ι*(*c*_*ij*_ = 1) = 5.2 bits and *ι*(*c*_*ij*_ = 0) = 0.04 bits. Core identity labeling is much more balanced with *ι*(*b*_*i*_ = 1) = 1.7 bits and *ι*(*b*_*i*_ = 0) = 0.6 bits for the target x-ray crystal structures.

We evaluate the efficiency of the encoded information as follows. First, pass a certain random fraction of the native labels 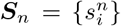 per target resulting in *N*_*r*_ total restraints. Construct the predicted labels ***S***_*p*_ for the same set of randomly selected residues for each target. Determine the Spearman correlation between *φ*(**S**_*n*_, **S**_*p*_) and LDDT over all targets. The total bits of information used per residue is then 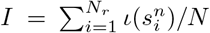. We repeat the random selections until *I* converges for each *N*_*r*_. *I* quantifies the amount of information used and *ρ* quantifies how useful that information is for predicting LDDT, i.e. encoding each protein structure. In Fig. 2, we plot *ρ* versus *I* for four orthogonal physical metrics **S**: C_*α*_ contact maps, core identity, secondary structure, and hydrogen-bond satisfaction.

**FIG. 2.**
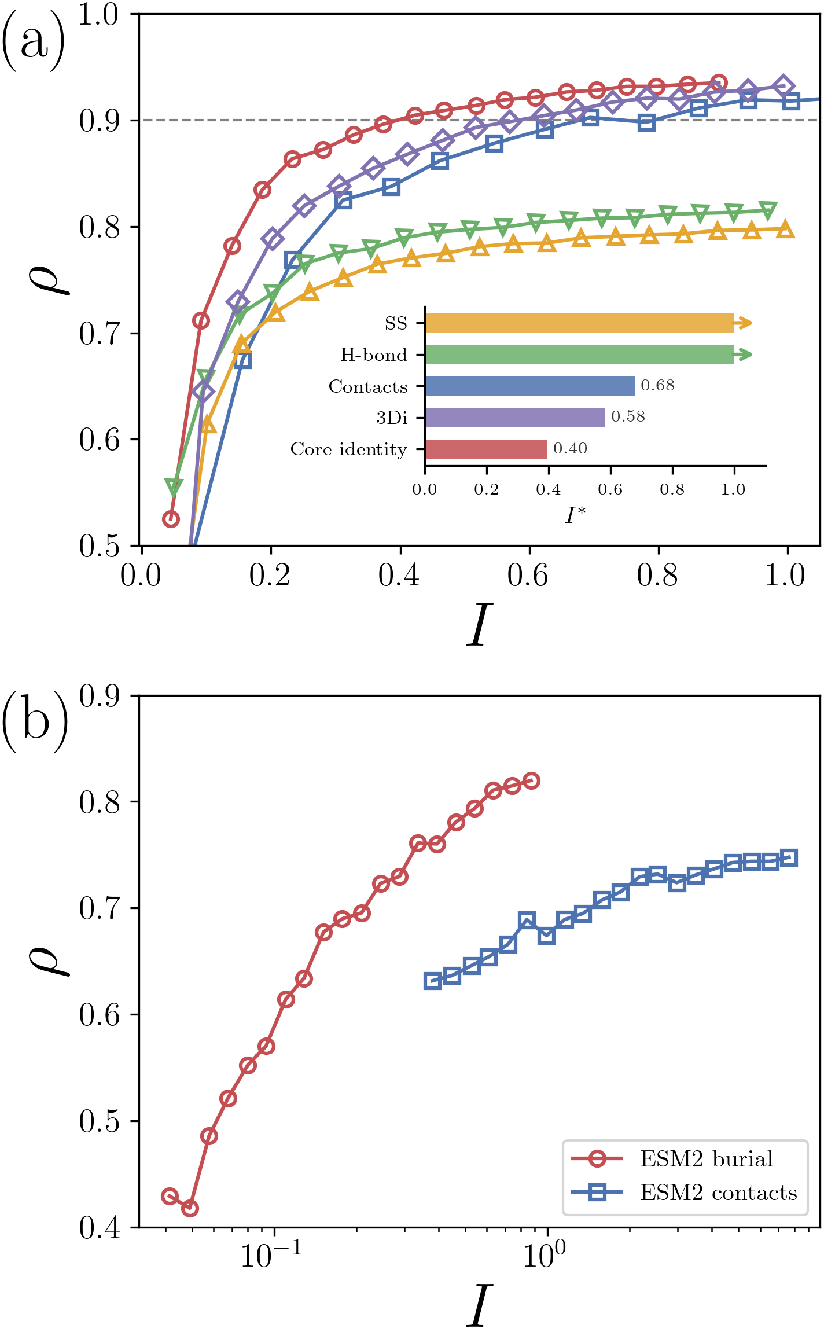
(a) The Spearman correlation *ρ* between LDDT and *φ*(***S***_*n*_, ***S***_*p*_) plotted versus the information content used *I* for the C_*α*_ contact maps (“Contacts”, blue squares), core identify (red circles), secondary structure (“SS”, yellow upward triangles), hydrogen-bonding satisfaction (“H-bond”, green downward triangles), and the 3Di embedding (purple diamonds). Inset: The amount of information *I*^*^ needed to obtain *ρ* = 0.9 for each physical metric and machine-learning embedding in panel (a). The arrows for secondary structure and hydrogenbond satisfaction indicate they do not reach *ρ* = 0.9. (b) *ρ* plotted versus *I* when using ESM2 embeddings to predict the contact maps (blue squares) and core identity (red circles) on a log_10_-scale.

First, we find that *ρ* does not increase above ∼0.8 when we include all *N* labels for the secondary structure and hydrogen-bond satisfaction, i.e. neither contain enough information to robustly predict LDDT. Next, we find that using all *N*(*N* − 1)/2 contact pair labels results in a *ρ* = 0.95, but the full contact maps use a large amount of information with *I* ∼ 25 bits per residue. We consider *ρ* = 0.9 sufficient for accurately folding a protein and refer to the amount of information to achieve this cutoff as *I*^*^, although our results are robust to the specific choice of the cutoff. Significantly less information from the contact maps is needed to achieve *ρ* = 0.9 with *I*^*^ = 0.68 bit per residue, which is equivalent to using ∼5% of the pairwise contact information. Remarkably, we find that information about the core identity outperforms contacts with *I*^*^ = 0.37 bits per residue.

The four metrics (contact maps, core identity, secondary structure, and hydrogen bonding satisfaction) are all physical representations of protein structure. Is there a machine-learned representation of protein structure that is more efficient than core identify? The protein structure comparison method FoldSeek was developed to condense a three-dimensional protein structure into a sequence of a single labels per residue from a learned alphabet of 20 letters using the autoencoder 3Di [20]. The *N* length representation is used to compare protein folds. While not explicitly designed to generate the most efficient embedding, 3Di provides an example of a state-of-the-art machine-learned encoding of 3D protein structures [20]. We show in Fig. 2 (a) that 3Di is more efficient than contact maps, but it is less efficient than the core identity, with *I*^*^ = 0.61 bits per residue.

When attempting to predict LDDT for proteins when we do not have access to the target structure to calculate **C**_*n*_ and **B**_*n*_, we can instead use the protein sequence to predict both **C**_*n*_ and **B**_*n*_. The efficiency of core identity in predicting LDDT suggests that we should achieve higher values for correlations with LDDT by predicting core identity over the contact maps. ESM2 represents the state-of-the-art in embedding a protein sequence into a numerical vector for each residue that represents the residue type and its sequence context. ESM2 was trained on coevolutionary data of protein sequences, without structures [25, 26]. However, ESM2 was found to accurately predict contact maps from sequence. In Fig. 2 (b), we plot *ρ* (from *φ*(**C**_*n*_, **C**_*p*_) versus LDDT) versus *I* using ESM2-predicted contacts from the sequence, resulting in *ρ* = 0.75 when using all of the contact information. To predict the core identity labels from ESM2, we developed a small but effective feed-forward network as discussed in the End Matter. We find that using the core identity labels predicted from the same ESM2 information results in *ρ* = 0.82, which is a significant increase in performance by switching from predicting contacts to predicting core identity. A combined score consisting of the average rank order of ESM2 contacts and core identity has the same correlation with LDDT as core identity alone. The efficiency of core identity suggests that structure prediction pipelines like ESMFold, which have already been applied to hundreds of millions of genomic sequences, could achieve higher accuracy by incorporating prediction of core identity rather than relying on contact-based representations [26]. We find that the state-of-the-art machine learning prediction of LDDT, AlphaFold3Score, has a correlation of *ρ* = 0.9 on the same dataset [18]. See End Matter for details.

When predicting **B**_*n*_ from sequence, there will be errors in the core identity labels, which will result in a lower performance in predicting LDDT. For example, the ESM2 prediction of **B**_*n*_ yields *ρ* = 0.82, whereas the perfect labeling results in *ρ* = 0.93. How sensitive is the correlation between *φ*(***B***_*n*_, ***B***_*p*_) and LDDT to errors in **B**_*n*_? Starting from the target structure’s true core identities, we randomly flip core identity labels with an increasing probability *p* resulting in a fraction of flipped labels *f*_flip_. Using the perturbed labels as a new **B**_*n*_ results in a new set of *φ* with reduced correlations with LDDT. In Fig. 3, we plot the LDDT correlation *ρ* versus *f*_flip_ with increasing *p* (black circles) and find that *φ* is robust to random errors. In particular, the correlation does not drop below *ρ* = 0.9 until *f*_flip_ ∼ 0.1.

**FIG. 3.**
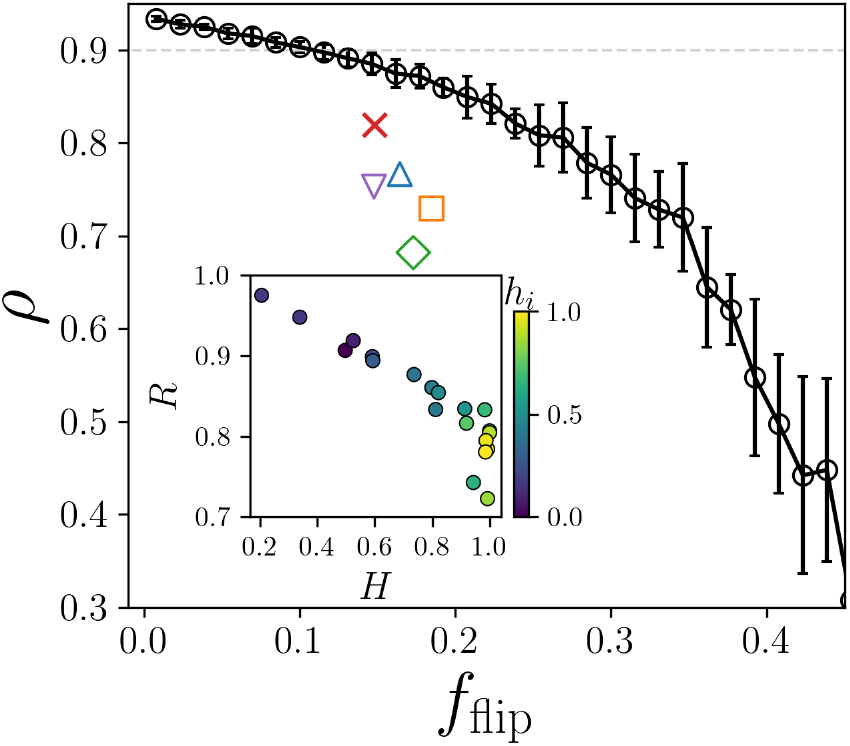
The Spearman correlation *ρ* between LDDT and *φ* plotted against the fraction of flipped core identity labels compared to the native labels. Errors were introduced randomly starting from the true identities (black circles and solid line). Error bars indicate *±* one standard deviation from the average over random samples. Core identity was also predicted from sequence using a series of models: the ESM2 model developed here (red x), Netsurfp3.0 (blue upward triangle), PaleAle6 (purple downward triangle), RaptorX (orange square), and E-pRSA (green diamond). The grey dotted line marks *ρ* = 0.9, our cutoff for a high quality predictor. Inset: The rate *R* of correct core identity labels plotted versus the entropy of the core identity labels *H* colored by hydrophobicity *h*_*i*_ (increasing from blue to yellow).

Next, we plot the LDDT correlation versus the fraction of flips from the correct labels for the ESM2 model, as well as additional methods that predict core identity from sequence: NetSurfP3.0 [27], PaleAle6 [28], RaptorX [29], and E-pRSA [30]. For each method, the target sequences are used to predict core identity, resulting in a fraction of flipped labels and a *φ* versus LDDT correlation. We find that all methods fall below random error, indicating that the residues that the core identity predictors miss are not random, but are instead the labels that are more important for predicting LDDT. Additionally, the ESM2 method achieves the highest correlation of *ρ* = 0.82 and is the closest to the random error.

To better understand on which residues the predictors fail, we investigate the accuracy rate *R* at which core residues are correctly labeled core and surface residues are correctly labeled surface for each residue type. Additionally, to quantify the base rates of surface and core are per residue type, we measure Shannon’s entropy *H* = −*p*_*c*_log_2_(*p*_*c*_) − *p*_*s*_log_2_(*p*_*s*_), for the probability that a residue type is in the core *p*_*c*_ versus on the surface *p*_*s*_. We indicate each residue type by its average hydrophobicity *h*_*i*_ from a consensus hydrophobicity scale [31]. In the inset of Fig. 3), we plot *R* versus *H* colored by hy-drophobicity for the ESM2 model and find that there is a strong linear negative correlation between how difficult a residue type’s core identity is to predict and the entropy of the base rates. While nearly all charged residues are on the surface (*H* ∼ 0), there is only a 50% chance for the most hydrophobic residues to be buried (*H* ∼ 1), and it is the hydrophobic residues whose core identity labels are the most difficult to predict. Retraining the ESM2 core identity prediction to focus on the hydrophobic residues did not result in meaningful improvement to performance. All core identity predictors tested show the same trend in *R* versus *H*.

What physical constraints determine which residues occur in the core? The prevailing hypothesis of protein folding is that hydrophobic collapse drives hydrophobic residues into the core and that the native core maximizes the number of hydrophobic core residues, while also respecting geometric constraints [4]. We quantify the core hydrophobicity as

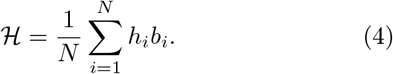

We compute the difference between the native and predicted core hydrophobicity Δℋ = ℋ_*n*_ − ℋ_*p*_. In Fig. 4, we plot the probability distributions *P*(Δ ℋ) for all predicted cores compared to the native cores binned by aver-age LDDT. While well-folded structures with high LDDT have a Δℋ ∼ ′, we find that ∼23% of predicted structures with the wrong fold (LDDT < 0.8) have a more hydrophobic core than the native fold (Δℋ < 0). We tested numerous hydrophobicity scales from the literature and found similar results to that in Fig. 4 [31–34]. Therefore, we find that hydrophobicity maximization using current hydrophobicity scales does not accurately determine core identity nor the native fold.

**FIG. 4.**
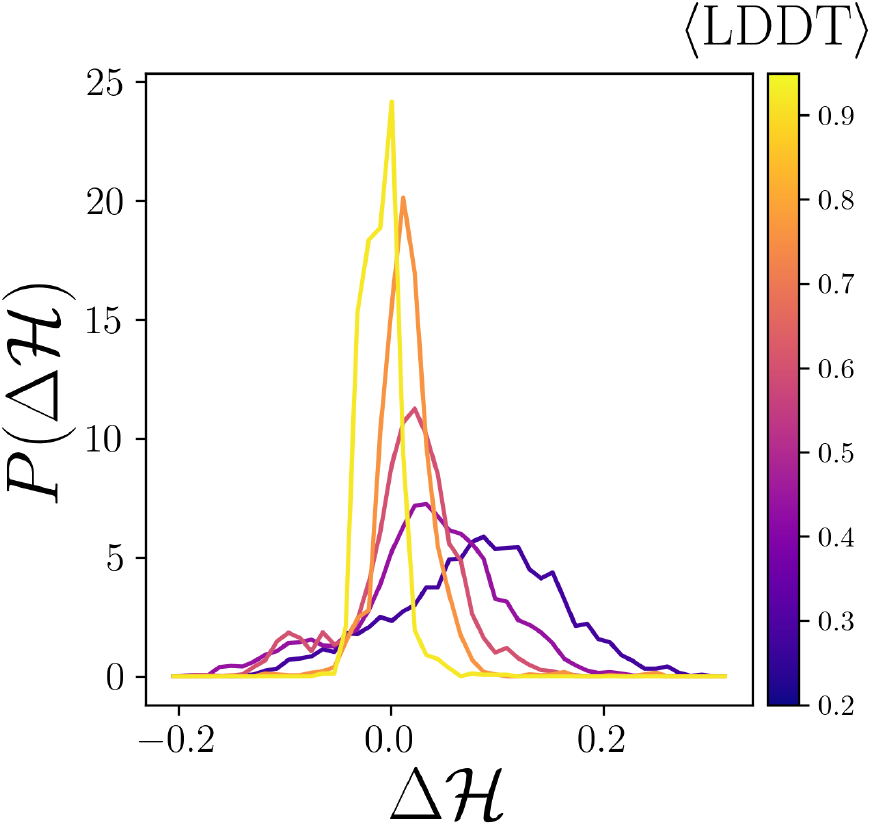
The probability distribution *P* (Δℋ) of the deviation of core hydrophobicity Δℋ = ℋ_*n*_ − ℋ_*p*_ binned by increasing LDDT (from purple to yellow).

In this Letter, we demonstrated that core identity, a binary label indicating which residues are core or surface, captures nearly all of the information necessary to encode the protein fold. Moreover, core identity is more efficient at this encoding than any physical or machine-learned metric that we tested. However, the simplicity of core identity labeling is deceptive. While the ESM2 based model achieves the highest accuracy predicting LDDT from sequence tested here, it does not achieve the *ρ* = 0.9 threshold for fully folding a protein from sequence. We find that the errors across core predictors occur predominantly for hydrophobic residues that contribute most to fold quality. That is, the residues that are hardest to predict are precisely those that matter most for achieving accurate folds. Additionally, we find that the current hydrophobicity scales cannot determine the native core from incorrect cores via maximization of ℋ.

A key aspect for future work will be to determine how well core identity can be used to guide molecular dynamics (MD) simulations of protein folding. Prior work has emphasized the benefits of pairwise contacts since they are simple to enforce using harmonic restraints. However, numerical spatial derivatives of SASA have been developed and can be used to enforce core identity during MD simulations [35, 36].

Crucially, these results reframe the protein folding problem. Rather than asking how the full sequence specifies protein structure, the central question can be reduced to: what determines the core identity of the subset of core residues that are difficult to predict? Whether the difficult-to-predict core residues reflect inadequacies in how we quantify hydrophobic packing or whether their burial is governed by factors beyond hydrophobicity remains an open and important question.

ATG acknowledges support from grant number 2023-329572 from the Chan Zuckerberg Initiative DAF, an advised fund of Silicon Valley Community Foundation. JS and CSO acknowledge NIH Training Grant No. T32GM145452. The authors acknowledge support from the High Performance Computing facilities operated by Yale’s Center for Research Computing.

## END MATTER

### Datasets

We used two datasets of protein structures for this work. First, we constructed a database of high resolution x-ray crystal structures of proteins that differ in sequence by more than 20% using the PDB culling server PISCES [37, 38]. This dataset includes 2,028 monomeric proteins with resolutions less than 1.8 Å. The matched protein target and predicted structure dataset was pulled from the protein folding competitions CASP11-15. We selected sets in which the target structure was determined using x-ray crystallography with a resolution less than 2.5 Å, resulting in 63 target structures and 24, 364 pre-dicted structures. CASP predicted structures often differ from their target structure’s sequence by a few residues on the ends. Therefore, before comparing rSASA values, we first aligned the sequences and trimmed the ends until the length of the target and predicted structure match exactly. To ensure no data leakage between the training set and CASP test set, we performed pairwise global sequence alignments between all 63 CASP targets and the 2,028 proteins in the PISCES training set using BLO-SUM62 with gap open and extension penalties of -10 and -0.5, respectively. No CASP target exceeded 42% identity with any protein in the training set.

We calculate rSASA using `FreeSASA` [24] with a probe size of 1.4 Å and a custom set of validated atomic radii [13, 39, 40]. Backbone secondary structure and hydrogen-bonding satisfaction were quantified using `DSSP` [41].

To account for the non-uniform distribution of LDDT values in the CASP dataset, all reported Spearman correlations use bootstrap resampling with a flattened LDDT distribution. LDDT is divided into 20 equally spaced bins and an equal number of samples are drawn with replacement from each bin. This process is repeated 10^3^ times and the median correlation is reported. Without flattening, the correlation is dominated by low accuracy structures and even using all contact map information will not yield large correlations with LDDT.

The binary Matthew’s correlation coefficient is defined in Eq. 2 in the main text. The 3Di embeddings and secondary structure features have more than two labels. Therefore, for these multi-label cases where there are *N*_*k*_ labels, we used a generalization of *φ*,

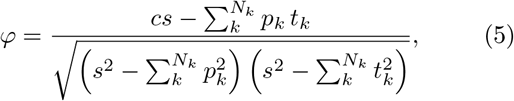

where s = Σ_*ij*_ *M*_*ij*_ is the total label count, *c* = tr(**M**) is the number of correct predictions, *p*_*k*_ = Σ_*i*_ *M*_*ik*_, and *t*_*k*_ = Σ_*j*_ *M*_*kj*_ [42].

A cutoff rSASA_*c*_ = 0.1 was used in this work to determine core identity. In Fig. 5, we show the Spearman correlation *ρ* between *φ* and LDDT as a function of rSASA_*c*_. We find that a wide range of cutoffs give similar values for *ρ*. We chose rSASA_*c*_ = 0.1 since it is the most strict definition of residue burial that still achieves the largest values of *ρ*.

**FIG. 5.**
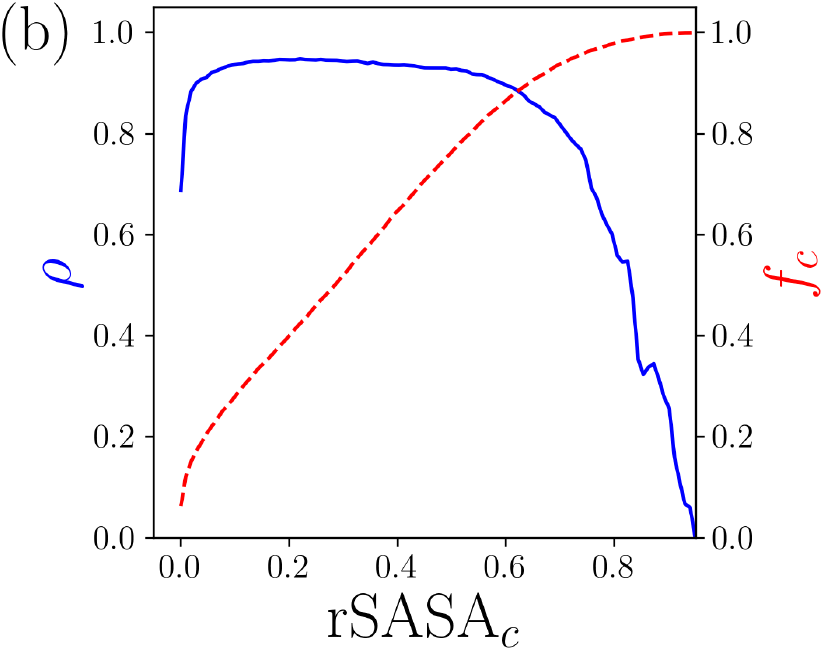
The Spearman correlation coefficient *ρ* between LDDT and *φ*(**C**_*n*_, **C**_*p*_) (blue line, left vertical axis) and the resulting fraction of core residues *f*_*c*_ (red dashed line, right vertical axis) plotted as a function of rSASA_*c*_.

### Core prediction

Residue core identity was predicted from sequence using four current methods in the same way that they were implemented on their respective servers at the time of publication. We also developed a simple ESM2-based core predictor. ESM2 takes a protein’s sequence as input and returns a high-dimensional vector for each residue that encodes both the residue identity and its context within the sequence. The length of the embedding is determined by the size of the ESM2 model that was implemented. We find the same results using the ESM2 650M and ESM2 3B parameters, and therefore we used the smaller model which outputs an embedding with length 1024 from the final layer 33.

We first embedded all of the sequences in the high-resolution x-ray crystal structure database. Then, we constructed a simple feed-forward neural network that takes each residue embedding and predicts the core identity of each residue. The network consists of two hidden layers, with size 256, ReLU activations, and dropout at 0.2. We use the Adam optimizer with a learning rate of 10^−3^, a batch size of 256, a 90/10 training/validation split at the protein level, and early stopping with a patience of 3 epochs. Training performance typically converged within 5 epochs.

We found no improvement using more complicated architectures with the same residue embeddings and when combining different layers (8, 16, 24 and 33) of ESM2 as input. Additionally, predicting rSASA as a regression task for downstream prediction of LDDT was as effective as classifying residue core identity. We also weighted the training to focus on predicting hydrophobic residues correctly, since our results indicate that they are more important for LDDT prediction, but the higher weighting of hydrophobic residues did not further improve the downstream LDDT prediction.

ESM2 contacts were predicted using the published attention-based model [26]. In that work, ESM2 contacts are defined using C_*β*_ atoms that are separated by 6 residues in sequence space and closer than 8 Å. We use the same definition when computing *φ*(***C***_*n*_, ***C***_*p*_) in Fig. 2 (b). We also weighted the contact map information by sequence separation, but found that neither close, medium, or long-range contacts in sequence were any better than using all contacts as a single type of label.

The code to predict core identity and then the average LDDT is available at: https://github.com/agrigas115/core_identity_score. The score requires a single CPU and approximately a minute of computation time per structure. The probability distribution of LDDT and the ESM2 predicted core identity score *φ*(***B***_ESM2_, ***B***_*p*_) is shown in Fig. 6 (a) and has a correlation of *ρ* = 0.82. The source code for AF3Score was obtained from https://github.com/Mingchenchen/AF3Score, which is used in its primary publication [18]. We ran AF3Score on NVIDIA L40S GPUs, using one GPU with 24 GB of memory and 16 CPUs with 80 GB of memory to score each structure in one minute of compute time. LDDT versus AF3Score is shown in Fig. 6 (b) and has a correlation of *ρ* = 0.9.

**FIG. 6.**
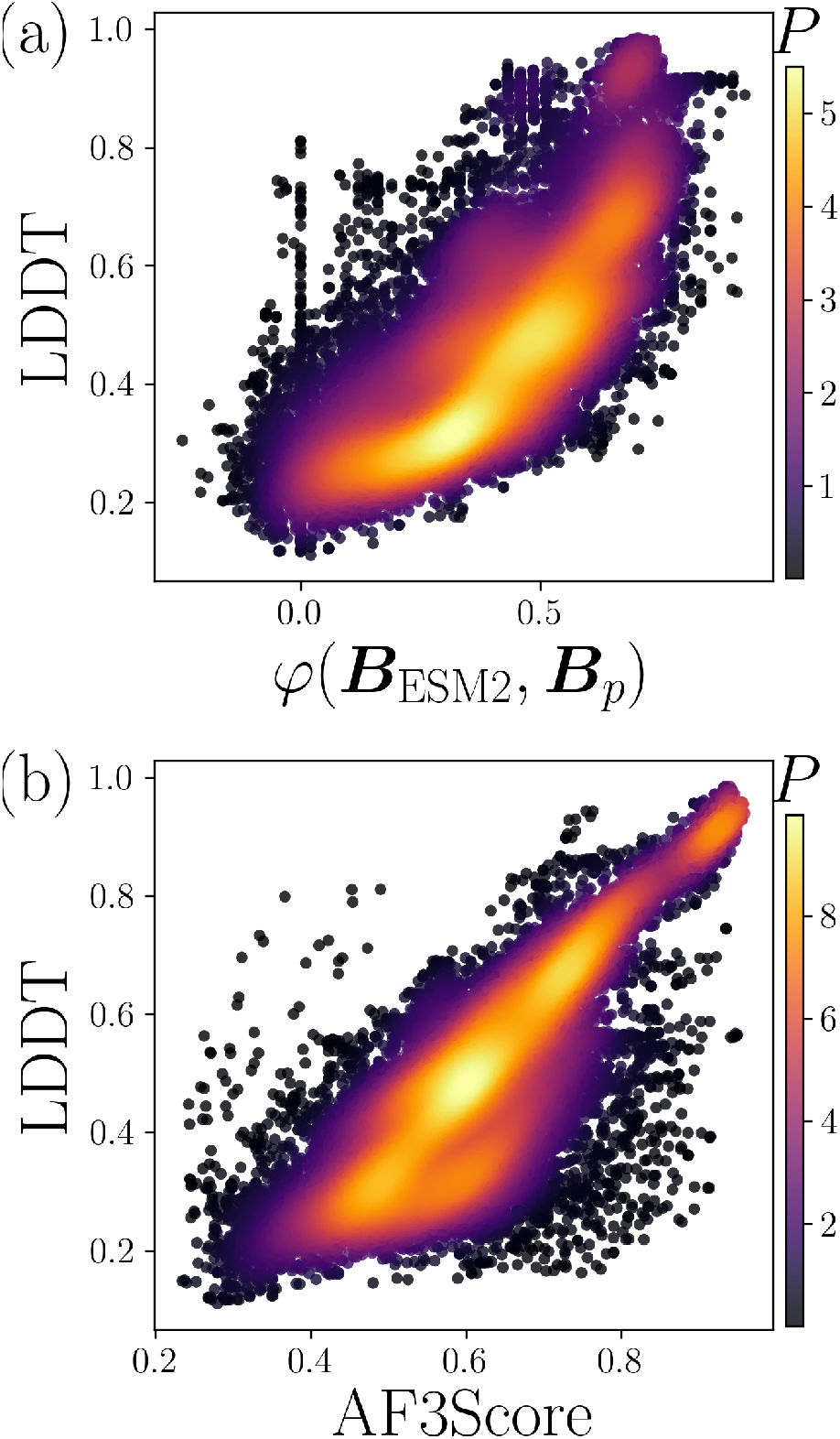
The probability density *P* (increasing from dark to light) of LDDT versus (a) the match in the core identity *φ*(***B***_ESM2_, ***B***_*p*_) when using ESM2 to predict the core identities ***B***_ESM2_ from sequence and (b) the AlphaFold3 Score (AF3Score).

